# Targeting Neuroplasticity to Improve Motor Recovery after Stroke

**DOI:** 10.1101/2020.09.09.284620

**Authors:** Sumner L. Norman, Jonathan R. Wolpaw, David J. Reinkensmeyer

## Abstract

After neurological injury, people develop abnormal patterns of neural activity that limit motor recovery. Traditional rehabilitation, which concentrates on practicing impaired skills, is seldom fully effective. New targeted neuroplasticity (TNP) protocols interact with the CNS to induce beneficial plasticity in key sites and thereby enable wider beneficial plasticity. They can complement traditional therapy and enhance recovery. However, their development and validation is difficult because many different TNP protocols are conceivable, and evaluating even one of them is lengthy, laborious, and expensive. Computational models can address this problem by triaging numerous candidate protocols rapidly and effectively. Animal and human empirical testing can then concentrate on the most promising ones. Here we simulate a neural network of corticospinal neurons that control motoneurons eliciting unilateral finger extension. We use this network to (1) study the mechanisms and patterns of cortical reorganization after a stroke, and (2) identify and parameterize a TNP protocol that improves recovery of extension force. After a simulated stroke, standard training produced abnormal bilateral cortical activation and suboptimal force recovery. To enhance recovery, we interdigitated standard trials with trials in which the teaching signal came from a targeted population of sub-optimized neurons. Targeting neurons in secondary motor areas on 5-20% of the total trials restored lateralized cortical activation and improved recovery of extension force. The results illuminate mechanisms underlying suboptimal cortical activity post-stroke; they enable identification and parameterization of the most promising TNP protocols. By providing initial guidance, computational models could facilitate and accelerate realization of new therapies that improve motor recovery.

## Introduction

Activity-dependent neuroplasticity occurs throughout life, affecting the CNS from cortex to spinal cord (Wolpaw and Carp, 2006; Wolpaw, 2010, 2018; Cramer *et al.*, 2011). When trauma or disease (e.g., stroke, spinal cord injury) impairs motor function, traditional rehabilitation concentrates on intensive practice of the impaired motor skills. Although this usually produces some recovery, significant disability often remains. Thus, the present challenge is to guide CNS plasticity so as to maximize functional recovery (S. C. Cramer et al., 2011; Dimyan and Cohen 2011; Wolpaw, 2012, 2018). Targeted-neuroplasticity (TNP) protocols are a new approach to addressing this challenge.

A TNP protocol creates a sensorimotor interaction with the CNS that induces activity-dependent plasticity at a key site (e.g., a particular spinal reflex pathway, a specific region in motor cortex) (Chen *et al.*, 2006; Buch *et al.*, 2008; Sitaram *et al.*, 2012; Ramos-Murguialday *et al.*, 2013; Norman *et al.*, 2018). This plasticity improves function. By doing so, the targeted plasticity enables activity that produces wider beneficial plasticity at other important sites (Wolpaw 2018). For example, after incomplete SCI, a TNP protocol that weakens a hyperactive spinal reflex can reduce the ankle clonus or foot-drop that prevents effective locomotor practice; it can thereby enable more effective practice, which produces wider beneficial plasticity (Thompson, Pomerantz, and Wolpaw 2013; Thompson and Wolpaw 2019).

While TNP protocols are a promising new therapeutic approach, their design and evaluation are formidable tasks. The many kinds of CNS plasticity, the many sites where they occur, the many new biological and technical methods, and the probability that the best therapies will combine several methods, generate an overwhelming number of appealing protocols. Testing even one of them is lengthy, demanding, and expensive, especially in humans. Computational models offer a solution. They can provide rapid and efficient screening of many potential protocols; only the most promising ones would then be tested in animals and/or humans (Reinkensmeyer *et al.*, 2016; Sedda *et al.*, 2018).

This study develops a neural network model of motor corticospinal plasticity before and after a simulated stroke and uses it to predict the therapeutic efficacy of different TNP training protocols. The results provide insight into the mechanisms of cortical reorganization after stroke and into the design of maximally beneficial TNP protocols. They indicate how judicious use of computational models might shape the development of effective new rehabilitation therapies.

## Materials and Methods

We simulated the impact of different treatment protocols on recovery of contralateral finger extension after a stroke that damaged motor cortex in one hemisphere. The foundations of this mathematical model are based on the structure and learning model first presented in Reinkensmeyer et al. (2012). That model investigated use-dependent recovery of movement strength following a stroke using a network of corticospinal (CS) neurons connected to downstream motor neuronal pools. The network learned using a biologically plausible reinforcement learning rule. To investigate the impact of different TNP protocols following a stroke, we now extend the structure of the model to represent CS neurons in both hemispheres; each neuron has its own connection strength to the motor neuronal pool *and* an intrinsic firing rate variability. We then simulate three scenarios: 1) the undamaged network underwent standard finger extension training; 2) the trained network was damaged by a stroke affecting contralateral motor cortex (i.e., contralateral to the finger) and then underwent standard finger extension training trials; and 3) the trained network was damaged by a stroke affecting contralateral motor cortex and underwent standard finger extension training trials interspersed with trials in which trial outcome was determined by the behavior of a specific population of CS neurons (i.e., TNP trials).

### Architecture

The model incorporates a network of *n* CS neurons that fire with activation levels *x_i_* (assumed to vary between 0 and 1 and to correspond, proportionally, to firing rates). Each CS neuron is connected to a motoneuronal (MN) pool via a scalar connection weight *w_i_*. A The MN pool sums the product of the neuron activation *x_i_* and connection weight *w_i_* using a saturation nonlinearity, *g_i_*.

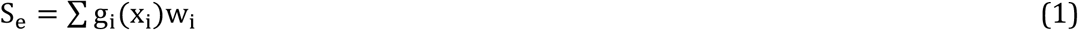

where function *g_i_* sets the saturation limit of neuron *i.* In this presentation, the model has a constant saturation limit of+1 for all neurons:

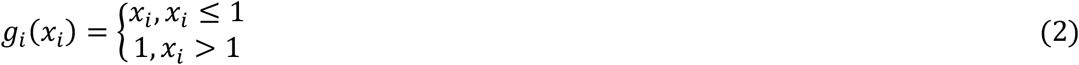

The MN pool activation level *S_e_* is proportional to a unitless finger extension force *F_e_*. Thus, the finger extension force generated is proportional to, and determined by, the weighted summed output of the CS network activation pattern. We found that varying the saturation limits across neurons did not significantly affect network dynamics. Thus, this presentation uses the constant saturation limit of +1 for all neurons.

### Network learning

The goal of the network is to learn the CS neuronal activation pattern that produces the maximum possible finger extension force, i.e., the best performance. Since the network consists only of neurons that excite extensor motor neuronal pools, the optimal activation pattern is achieved when the activation of every neuron is increased to the neuron’s saturation limit of +1.

To learn this pattern, the network employs an iterative reinforcement learning protocol: after each trial (i.e., each movement attempt) the network adjusts the activation patterns based on a scalar teaching signal, which is finger extension force *F_e_*. Using a single signal to optimize a large network presents a credit assignment problem: if finger extension force *F_e_* increases on a given trial, which neurons are responsible for the increase? Reinforcement learning can solve this credit assignment problem, albeit imperfectly (Mazzoni *et al.*, 1991; Williams, 1992; Anderson *et al.*, 1997; Werfel *et al.*, 2005; Reinkensmeyer *et al.*, 2012). As will become clear below, TNP trials are designed to mitigate the credit-assignment problem and to thereby improve the learning outcome.

We implement reinforcement learning with stochastic search. The algorithm uses a noise process to generate a new activation pattern for each trial (i.e., each attempted movement). If the new activation pattern increases finger force compared to the previous trial, the algorithm stores (i.e., switches to) the new activation pattern. The stochastic search algorithm is a simplified form of the random search with chemotaxis algorithm (Anderson *et al.*, 1997):

Given an initial activation pattern *X_0_* that produces a force *F_0_*:

1. Activate CS neurons with pattern *X_i_*=*X_0_*+*v_i_*, where *v_i_* is random noise, and measure the force *F_i_* produced by this pattern.
2. Store (i.e., switch to) the new pattern *X_i_* if the force *F_i_* it produces is greater than *F_0_* (i.e., if *F_i_*>*F_0_*, then let *X_0_*=*X_i_* and *F_0_*=*F_i_*).
3. Repeat

We also tested a gradient descent stochastic search method, another biologically plausible solution to the credit assignment problem (Werfel *et al.*, 2005). It produced comparable results; thus, we do not detail them here.

### Neuron parameters

The CS neurons in the model are described by their current activity level (*x_i_*, proportional to firing rate), and their excitatory synaptic connectivity to the extensor MN pool (*w_i_*). In the present work, we augment the model of (Reinkensmeyer *et al.*, 2012) to allow different levels of trial-to-trial firing rate variability for different CS neurons (*σ*). This feature is based on evidence (Faisal *et al.*, 2008) that neuronal variability differs across cortical areas. The network updates each neuron’s activation level after a successful movement trial (i.e., a trial that produces more force than the previous trial). Each neuron’s connectivity and variability remain constant throughout the simulation. In reality, spared descending pathways are plastic after an injury. Although we do not change weighting (*w_i_*), altering the firing rate is mathematically equivalent in this model; it is a heuristic for the dynamic nature of downstream synaptic plasticity.

Neuronal parameters, including initial activation level, fixed connectivity, and fixed variability, were initialized by sampling from lognormal distributions. Functional and structural parameters in the brain, including synaptic connectivity and firing rates, are typically not normally distributed; they are skewed with a heavy tail. Thus, they are closely approximated by lognormal distributions (Buzsáki and Mizuseki, 2014)(See Appendix, Neuron parameter distributions). To represent different cortical areas, we used different lognormal distributions (Figure 2).

### Synaptic connectivity

Monosynaptic and multisynaptic CS pathways are represented in the model by a single, fixed connectivity from each CS neuron to the finger extensor MN pool (i.e., weights labeled *w_i_* in Fig. 1). We explicitly represent both cortical hemispheres in the model, since the hemisphere ipsilateral to the moving finger is known to be able to activate the requisite MN pools through uncrossed pathways, and these pathways are thought to play a significant functional role after stroke (Cramer *et al.*, 1997). We designed the distributions of the neuronal connectivities to reflect known physiology: neurons from the hemisphere contralateral to the motor task are, on average, more strongly connected to the MN pools than ipsilateral neurons (Fig. 2) (Nudo *et al.*, 1992). Furthermore, these contralateral neurons outnumber the ipsilateral neurons by a ratio of 9:1. This is consistent with the physiological situation wherein about 90% of axons cross over from the lateral corticospinal tract and about 10% of fibers travel within the uncrossed anterior corticospinal tract (Martin and Jessell, 1996).

**Fig. 1.**
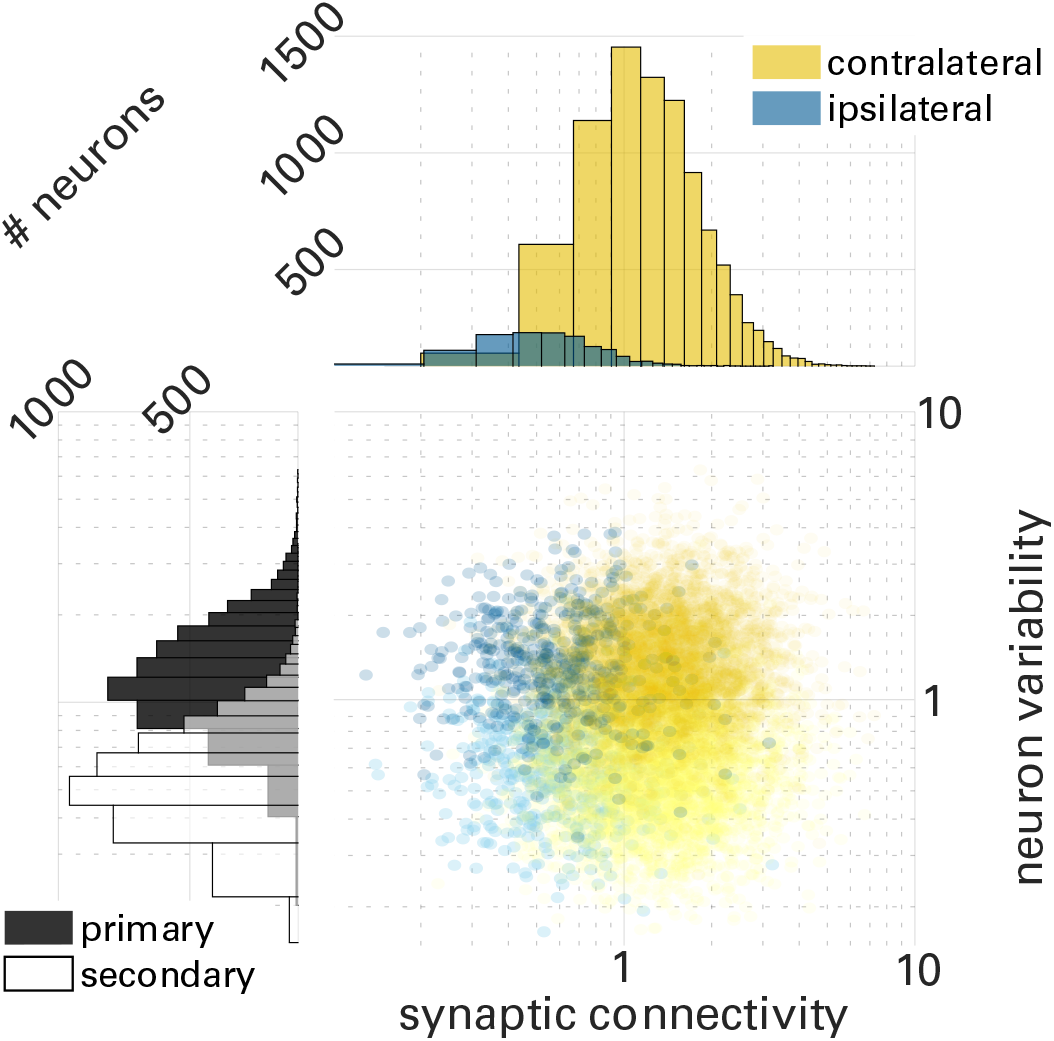
Network architecture. A two-layer feedforward neural network incorporates n corticospinal (CS) neurons with activation levels *x_i_*. These activation levels are generated when the neurons are given a command to maximize finger extension force *F_e_*. A motoneuronal pool *S_e_* sums the weighted activation pattern. A nonlinear function *g_i_* implements the physiological observation that the contribution of any single CS neuron to the excitation of the motoneuronal pool saturates at some activation level. The network optimizes the activation pattern *X* through reinforcement learning in simulated consecutive movement practice trials in which extension force *F_e_* is the teaching signal.

**Fig. 2.**
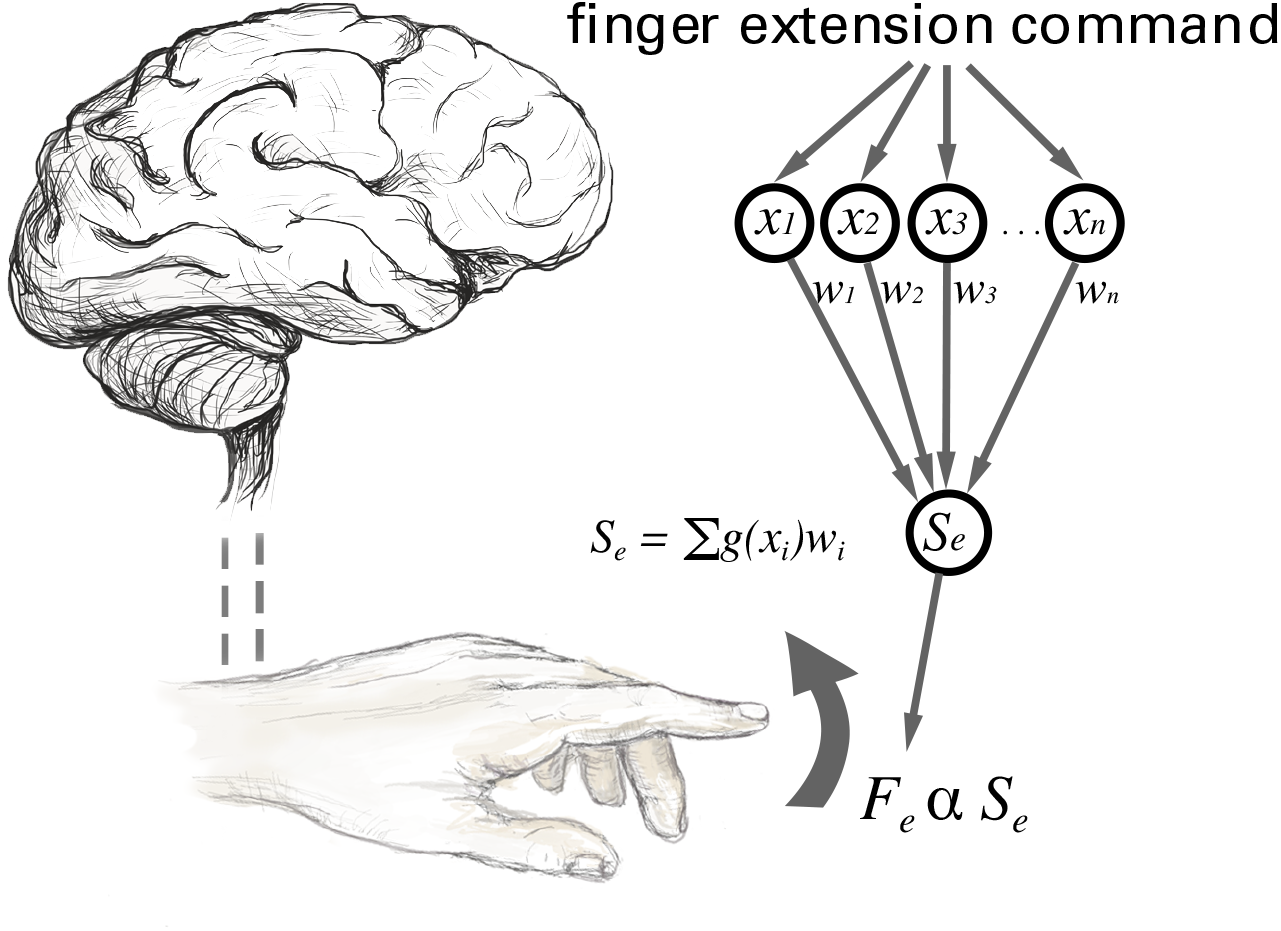
Parameter distributions for a network of 10,000 corticospinal (CS) neurons. Synaptic connectivity adheres to a bimodal distribution resulting from use of two lognormal probability density functions, one for the contralateral cortex, and one for the ipsilateral cortex. The mean of the bimodal distribution was chosen to be one. Most (90%) neurons reside in the cortex contralateral to the finger to be extended and have stronger connectivity (contralateral = yellow). The remaining neurons reside in the ipsilateral cortex and have weaker connectivity (ipsilateral = blue). Neuronal firing variability also adheres to a bimodal distribution arising from two lognormal probability density functions. Neurons in primary motor areas are more task-related and exhibit more trial-to-trial variability during movement attempts (primary = dark). Neurons in secondary motor areas are less task-related and exhibit less trial-to-trial variability during movement attempts (secondary = light). The resulting network has four broad types of neurons: 1, high-connectivity/high-variability (dark yellow); 2, high-connectivity/low-variability (light yellow); 3, low-connectivity/high-variability (dark blue); 4, low-connectivity/low-variability (light blue).

### Variability

Motor variability is necessary for motor learning (Herzfeld and Shadmehr, 2014; Wu *et al.*, 2014); higher levels of motor variability accompany motor skill acquisition. Furthermore, such variability is present from the neuronal level to the behavioral level (Faisal *et al.*, 2008). Thus, the algorithm incorporates trial-to-trial corticomotor variability to drive motor learning in our training scenarios.

The algorithm varies neuronal firing rate by sampling an activation noise level from a normal distribution that is specific to each neuron. A higher-variability neuron has a wider normal distribution; it averages a larger stochastic perturbation to its activation level on each trial compared to a lower-variability neuron. The most task-relevant brain areas exhibit more variability. Thus, the model gives neurons in primary motor areas of both hemispheres higher variability for the finger extension task than neurons in secondary motor areas. At the same time, high- and low-variability neurons are unlikely to be wholly separated spatially (van Steveninck *et al.*, 1997; Warzecha and Egelhaaf, 1999). For this reason, the model overlaps high- and low-variability neurons to some degree (Fig. 2). A neuron’s firing rate is related to its mean firing rate by a power function, the parameters of which vary across cortical areas and behavioral conditions (Dean, 1981; Lee *et al.*, 1998). Our model simplifies this relationship to clarify the effects of neuronal variability on the network dynamics: a neuron’s trial-to-trial variability differs across cortical areas but remains constant over time and does not depend on its firing rate.

### Simulations

We use this model to simulate three scenarios for learning finger extension: 1) learning by the uninjured network; 2) learning by the network following a unilateral stroke; 3) learning by the network following a unilateral stroke with TNP training trials interdigitated. The initial, uninjured network consisted of 1,000 CS neurons. To simulate the stroke, Scenarios 2 and 3 fix the activation and connectivity of CS neurons lost to the stroke to zero. To simulate the disruptive effects of the stroke on the properties of surviving CS neurons (Nudo and Milliken, 1996), their initial post-stroke activation patterns for Scenarios 2 and 3 are randomized.

Since network learning is driven by movement attempts (i.e., trials), the number (i.e., dosage) of movement trials affects the results. Lang et al. (2009) found that participants with stroke completed an average of 32 functionally oriented movements/day during upper extremity rehabilitation sessions. However, recent rehabilitation interventions have successfully administered 150-250 trials/session, for up to 23 sessions, for a total of 3450-5750 trials (Buch *et al.*, 2008). In the simulations presented here, the network trained in a given scenario for a total of 20,000 trials. We gave the same dosage of 20,000 trials in each training scenario to facilitate comparing their results.

### Scenario I: Learning with an undamaged network

To provide a baseline, we simulated learning in the uninjured network. For each trial, finger extension force was determined using all 1,000 neurons. The teaching signal was the difference from the previous trial in finger extension force. If the force was greater than that of the previous trial, the network switched to the new activation pattern.

### Scenario II: Learning after stroke

To simulate neuronal death after a stroke, we disconnected (de-weighted) a subpopulation of CS neurons by permanently setting their connectivity and activation levels to zero. This presentation focuses on the most severe stroke: a stroke that damages contralateral primary motor cortex, where high-connectivity, high-variability CS neurons are concentrated. The teaching signal from each trial is the finger extension force produced by the remaining intact CS neurons of both hemispheres.

### Scenario III: Learning after stroke with targeted neuroplasticity (TNP) trials interdigitated

Finally, we simulated learning the finger extension task after stroke with a portion of the training trials now dedicated to a TNP protocol. In TNP trials, only the targeted neurons determined the teaching signal. That is, in a TNP trial, separate training signals were generated for the total network (finger extension force, *I_A_*) and the targeted network subpopulation (targeted intervention, *I_B_*). The training algorithm used *I_B_* to determine success or failure (i.e., to determine whether the network switched to the new activation pattern). If *I_B_* indicated success, the parameters of all the surviving CS neurons (targeted and untargeted) were updated. The total number of trials did not change; Standard trials and TNP trials were interspersed in different ratios to determine the optimal dosage. Thus, we systematically evaluated the dependence of force recovery on two training parameters: 1) which subpopulation of CS neurons (defined by neuronal parameters and/or location) was targeted; and 2) the proportion of the total trials that were TNP trials (i.e., dosage).

## Results

We trained the network in three Scenarios: 1) Learning with an undamaged network; 2) Learning after stroke; and 3) Learning after stroke with targeted neuroplasticity (TNP) trials interdigitated. We ran each scenario ten times, with 20,000 trials each time. In Scenario 1, the undamaged network achieved 84.5% of the maximum possible finger extension force (i.e., the maximum force possible for all CS neurons). For simplicity, we scale all results to the average maximum force produced in scenario 1 (Fig. 3).

**Fig. 3.**
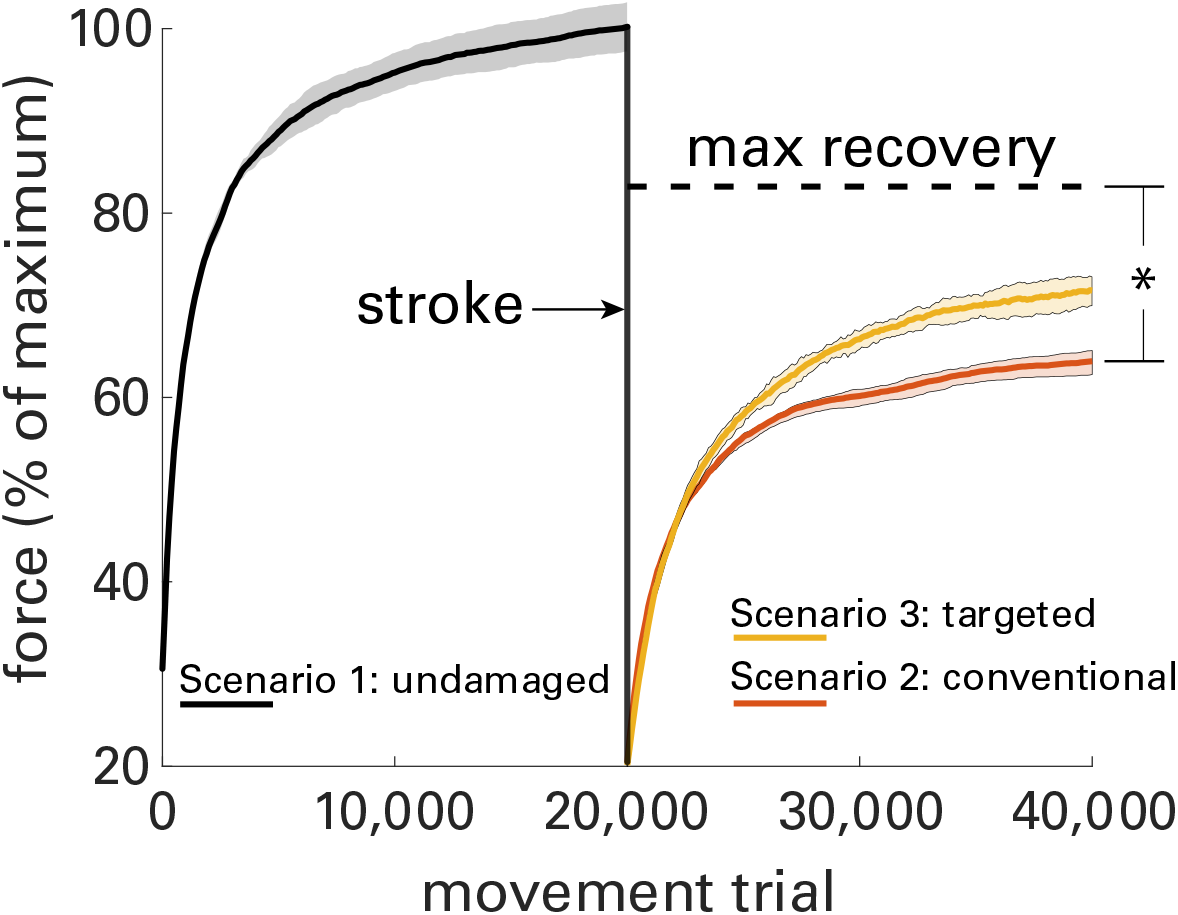
Force production by the three Training Scenarios. Force (in % of the maximum force achieved by the uninjured network after 20,000 trials) as a function of the number of movement trials. After a stroke, max recovery is the theoretical maximum force possible for the surviving CS neurons. Solid lines represent the mean result and shaded areas represent the standard deviation of ten simulations. Conventional Training (Scenario 2) leaves substantial residual capacity (*). Targeted plasticity (TNP) (Scenario 3) recovers much of this capacity; thus, it produces greater force recovery.

In Scenario 2, we simulated neuronal death due to stroke by removing a subpopulation of CS neurons from the network (by fixing their connectivities and activation levels at zero). We randomized the initial firing rates of the surviving CS neurons. The teaching signal was the finger extension force produced by all the surviving CS neurons. Finger force increased exponentially and then approached a recovery plateau in which it generated 64.9% (+/− 2.9% SD) of the maximum force possible before the stroke (Fig. 3, Scenario 2). While this is a substantial recovery, significant capacity to generate force remained unused (Fig. 3, residual capacity for recovery).

Scenario 3 began in the same way as Scenario 2, i.e. with a stroke and 20,000 trials of subsequent training. In our initial evaluation of Scenario 3, every fifth trial was a TNP trial; in which the teaching signal was provided by the surviving CS neurons in the secondary motor areas of both hemispheres. After this training period, the network reached 72.8% of the maximum force possible before the stroke, an increase over Scenario 2 (conventional rehab). Scenario 3 enabled the network to use 43.6% of the latent capacity for recovery that Scenario 2 did not capture.

## Impact of training on different neuronal types

Scenario 1 (the uninjured network) preferentially optimized activation of high-variability (M1, primary motor cortex), high-connectivity (contralateral) neurons. Thus, it left some residual capacity unachieved.

Scenario 2 (the injured network with standard training) also favored optimization of high-variability, high-connectivity neurons (Fig. 4c). However, because many such neurons were gone, the training then optimized high-variability neurons with low connectivity; it shifted activity toward the weakly connected, but undamaged ipsilateral hemisphere.

**Fig. 4.**
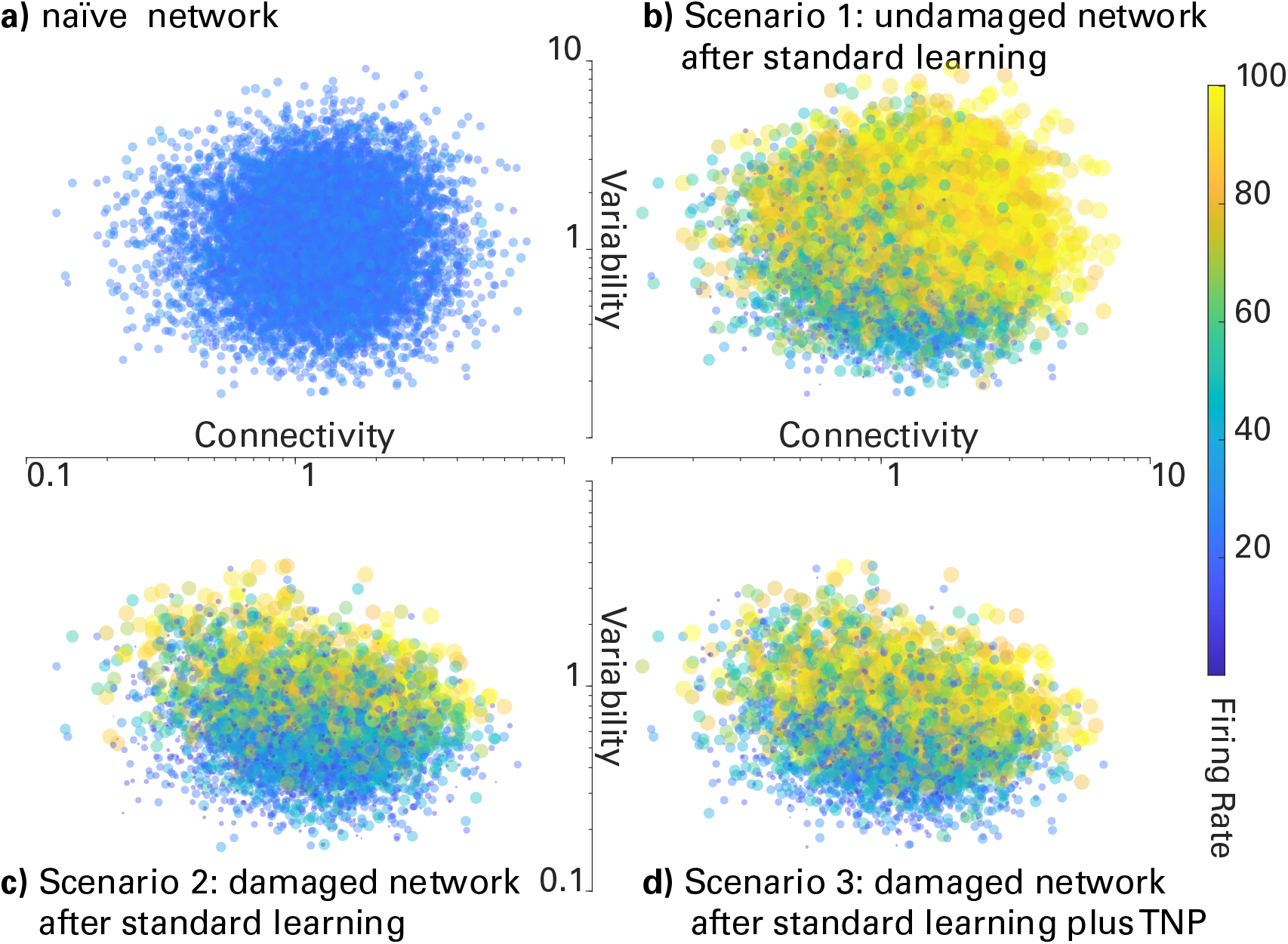
Neuronal activation patterns for the naïve network and for the network at the end of each of the three Scenarios (shown for a 10,000-neuron network). **a**, Normal network before learning. Network activation reflects a pseudorandom sample of activation levels. **b**, Normal network after standard learning and before stroke, with optimized neurons in yellow (high firing rate). High-connectivity/high-variability neurons are optimized. **c**, Injured network (i.e., 8125 remaining neurons) after standard learning. Optimization is focused on the remaining high-variability neurons; low-variability neurons are generally not optimized. **d**, Injured network after standard learning plus targeted neural plasticity (TNP). Optimization of low-variability neurons is increased (i.e., d is substantially more yellow than c). The result is that force recovery improves (Fig. 3).

Scenario 3 was the same as Scenario 2 except that every fifth trial based its outcome only on neurons in secondary motor areas of either hemisphere (i.e., generally low-variability neurons not optimized by Scenario 2). These targeted areas had the potential to significantly improve overall network performance. However, without TNP, the network struggled to access them. In standard trials, the stronger impact on trial outcome (i.e., success or failure) of the random changes in high-variability neurons masked the weaker impact of low-variability neurons. The TNP trials of Scenario 3 removed this masking. They promoted activation of low-variability/high-connectivity neurons (Fig. 4d).

## Impact of training on the topography of neuronal activation

To visualize the predicted effect of the three scenarios on the topography of the task-related neuronal activation, we mapped neuronal parameters to brain areas. Specifically, we mapped high-and low-connectivity neurons contralateral and ipsilateral to the movement. We mapped high and low variability neurons to primary motor cortex (M1) and dorsal premotor cortex (dPM; a secondary motor area), respectively (see Table 1). In Figure 5, we use this mapping to show network reorganization across hemispheres during learning before a stroke (Scenario 1), and after a stroke without (Scenario 2) or with (Scenario 3) TNP training.

**Fig. 5.**
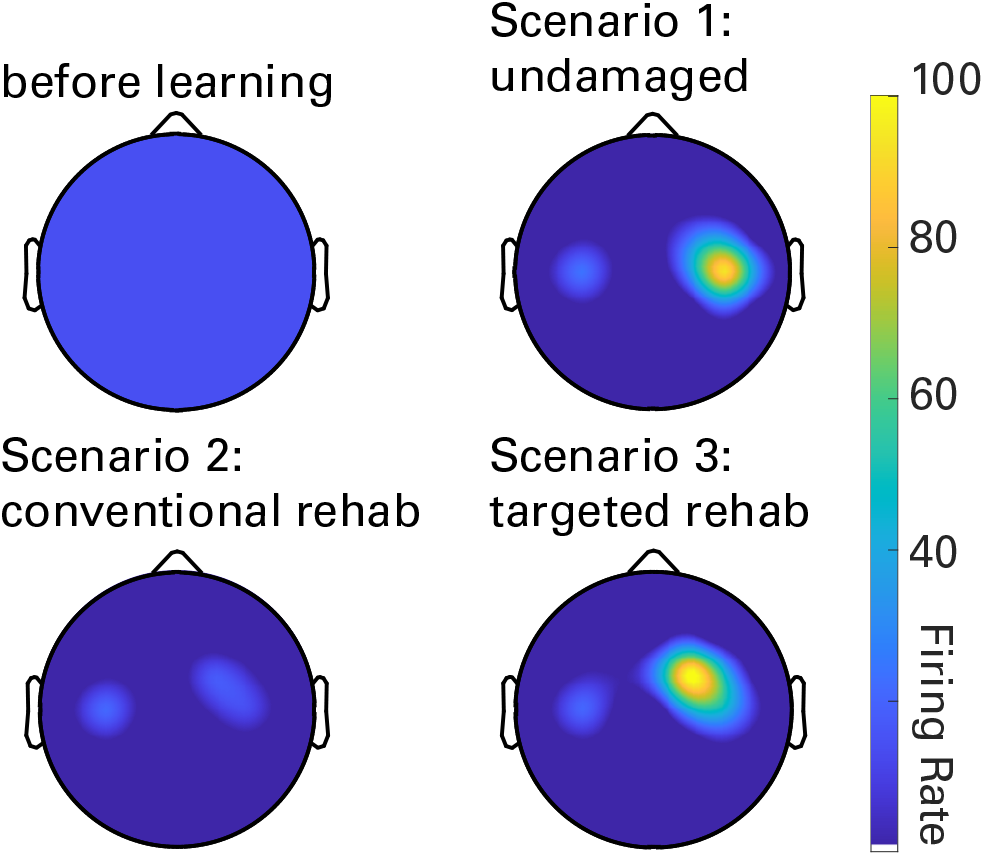
Topography of task-related neuronal activation before training and at the end of training for each of the three training Scenarios. **1**) standard training of the uninjured network; **2**, standard training of the injured network; and **3**, standard training of the injured network plus TNP. Scenario 1 primarily optimizes neurons in contralateral primary motor cortex (high connectivity/high variability neurons). Scenario 2 also optimizes neurons in ipsilateral (undamaged) cortex (high-variability/low-connectivity neurons). This produces abnormal bilateral activation and diffuse activation in the remaining contralateral (damaged) primary motor cortex. Scenario 3 also optimizes neurons in secondary motor areas (e.g., dPM); and it restores more normal laterality.

**Table 1:** Cortical Mapping of Neuronal Parameters (M1: primary motor cortex; dPM: dorsal premotor cortex)

|  |  |  |  |
| --- | --- | --- | --- |
| *variability* | *high* | *Ipsilateral-M1* | *Contralateral-M1* |
|  | *low* | *Ipsilateral-dPM* | *Contralateral dPM* |
|  |  | *low* | *high* |
|  |  | *connectivity* |  |

Standard learning by the intact network (Scenario 1) optimized contralateral primary motor (i.e., high-variability, high-connectivity) neurons. This resulted in contralateral activity (Fig. 5). Standard learning after stroke (Scenario 2) shifted activation to the undamaged ipsilateral hemisphere (i.e. toward high-variability neurons with relatively low connectivity). This produced bilateral activation for the unilateral task. Standard learning after stroke plus TNP (Scenario 3) also optimized neurons in secondary motor areas of both hemispheres (i.e., low-variability neurons). This produced more normal lateralized activation and improved force recovery.

## Optimizing target population and dose

One of the model’s advantages is its ability to reveal which populations to target to maximize therapeutic effect. To this end, we targeted different cortical areas and gave TNP on every fifth trial. We ran each simulation twenty times; each one simulated a stroke that affected a random subset of 75% of the CS neurons in contralateral primary motor cortex (i.e., mostly high connectivity, high variability neurons). We then determined for each area the improvement in residual force capacity recovered over that provided by standard training alone (Scenario 2) (Fig. 6a). Delivering TNP therapy to secondary motor areas (e.g. dPM) gave the best results, restoring 30.5% of the residual capacity left by Scenario 2. Targeting dPM only in the damaged hemisphere had similar effects. Targeting dPM in the undamaged hemisphere was ineffective. Targeting primary motor cortex (M1) either bilaterally or only in the damaged hemisphere caused maladaptive plasticity that reduced force production compared to Scenario 2. Targeting primary motor cortex only in the undamaged hemisphere had no significant effect.

**Fig. 6.**
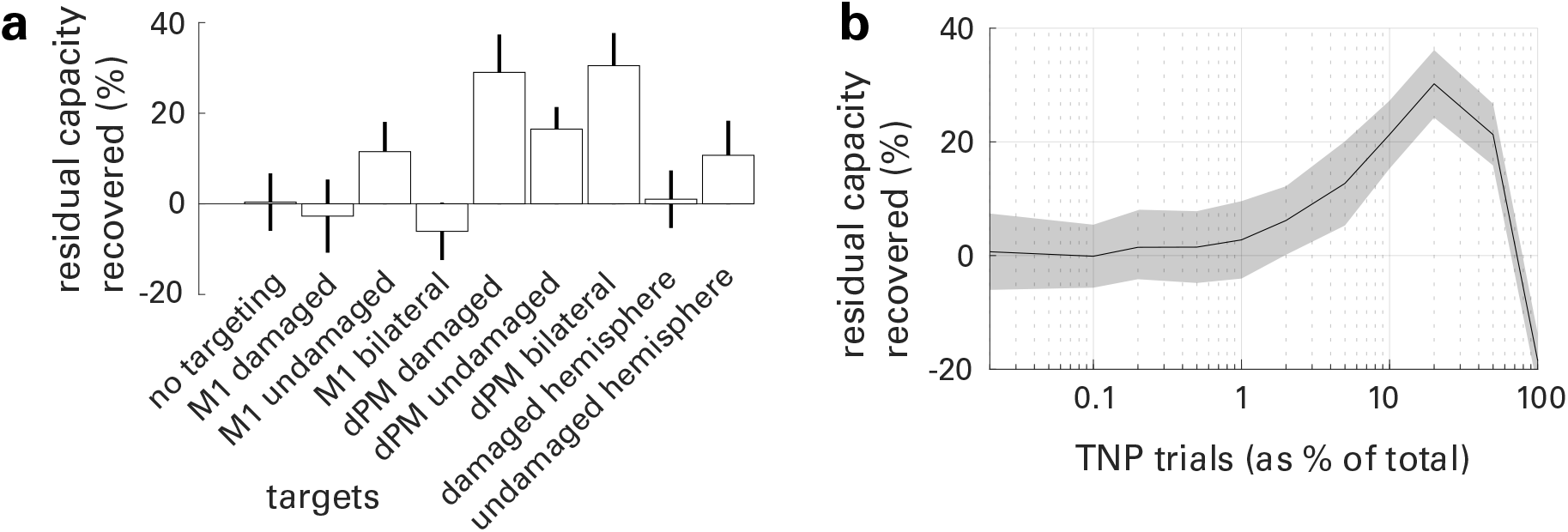
The benefits of targeted neuroplasticity (TNP): the impact of the area targeted and the TNP dosage. **a**, Additional residual force recovered by standard learning plus TNP (given on 20% of trials) over that recovered with standard learning alone, for each of eight different cortical regions (M1: primary motor cortex; dPM: dorsal premotor cortex). Results are given as a percentage of the residual capacity for recovery left uncaptured by Scenario 2 (i.e., no TNP trials). Positive values indicate a better outcome with TNP. Error bars are standard deviation of 20 simulations. **b**, Residual force recovery as a function of the percentage of trials that were TNP trials (results shown for targeting dPM). Solid line is the mean and shaded area indicates the standard deviation of 20 simulations. Recovery improved with increased dosage of targeted feedback. Recovery reached a maximum when 10% of trials were given targeted feedback and declined thereafter.

We then focused on the most beneficial targeting (i.e., dPM bilaterally) and assessed the effect of varying TNP dosage from 1/2000 trials to every trial. We ran each simulation twenty times; each one simulated a stroke that affected a random subset of 75% of the CS neurons in contralateral primary motor cortex (i.e., mostly high connectivity, high variability neurons). We then determined for each TNP dosage the residual capacity recovered compared to conventional rehabilitation (Fig. 6b). Doses <1% were ineffective. As dose increased from zero, training efficacy increased; it reached a maximum when 20% of the trials were TNP trials. Efficacy declined at doses >50%. The results were similar when other cortical areas were targeted. To our knowledge, these results provide the first substantive insight into the most effective dosage of TNP as a fraction of total training. The consistency of the result across different targeted areas suggests that the result may apply across different lesion types and training protocols.

## Mechanisms: Neuron-specific optimization rates and ‘blocking’

Why did activity lateralize in the undamaged brain, but become bilateral after the simulated stroke? And why did TNP help remediate this situation, improving force recovery? Two mechanisms account for these results. First, high-variability neurons optimize faster than low-variability neurons. Second, once optimized, high-variability neurons prevent (‘block’) optimization of low-variability neurons.

To illustrate these mechanisms, we used a Monte Carlo method wherein we ran the same, single trial of the model 10,000 times - once at the beginning of network training, and once after the network had been trained for 20,000 trials, when the network had learned to generate more force (Fig. 8a) by increasing activation across neuron groups (Fig. 8b). Using the results from the Monte Carlo simulations, we estimated the probability that each neuron group would, through their summed activity that resulted from the random perturbation on that trial, contribute toward a positive change in force. Specifically, we calculated the mean change in activation 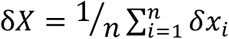 and the mean change in force 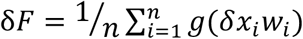 across all neurons *x_i_*=1,2,…,*n* in each group. We report these values for successful trials that advance the optimization (Fig. 8c-e) and for all trials (Fig. 8f-h).

## Highly variable neurons optimize first

Before training, no neurons are optimized (Fig. 8b). Thus, on learning trial 1, all neurons increase or decrease their activity with equal (i.e. 0.5) probability on each trial (Fig. 8f). Recall that we ran this first learning trial many times. Across all of these first learning trials that are successful (i.e., produce an increase in force), high variability (i.e., fast) neurons change activation (Fig. 8d), and thus their contributions to total force (Fig. 8e), by relatively large amounts. The contribution to increase in force is particularly high for the fast, high-connectivity (i.e., strong) neurons, which will cause them to optimize quickly. These neurons will saturate as training proceeds (they cannot exceed the maximum firing rate defined by the *g* function). At trial 20,000, they do not contribute any more to increasing the total force (Fig. 8e, after training), or it is extremely unlikely that they will do so (Fig 8c, after training). The network then favors at this later time stage (i.e., following the optimization of fast/strong neurons) the optimization of high-variability/low-connectivity (fast/weak) neurons (Fig. 8d, after training).

## The network learns at increasingly slow rates, failing to optimize some neurons

Saturated neurons cannot contribute to further increases in force production. Thus, for saturated neurons, the probability that δx is positive approaches zero (Fig. 8f). As observed above, more variable and strongly connected (fast/strong) neurons optimize more quickly, and thus, on average, saturate first. For the network to switch to a new activation pattern, yet-to-be optimized neurons must not only produce their own net positive effect on force but must also overcome any negative effect of the more-variable neurons that, once saturated, tend to decrease force on any given trial (Fig. 8e). This blocking phenomenon, created by the nonlinear saturation function, *g*, causes the latent residual capacity shown in Figure 3.

**Fig. 8.**
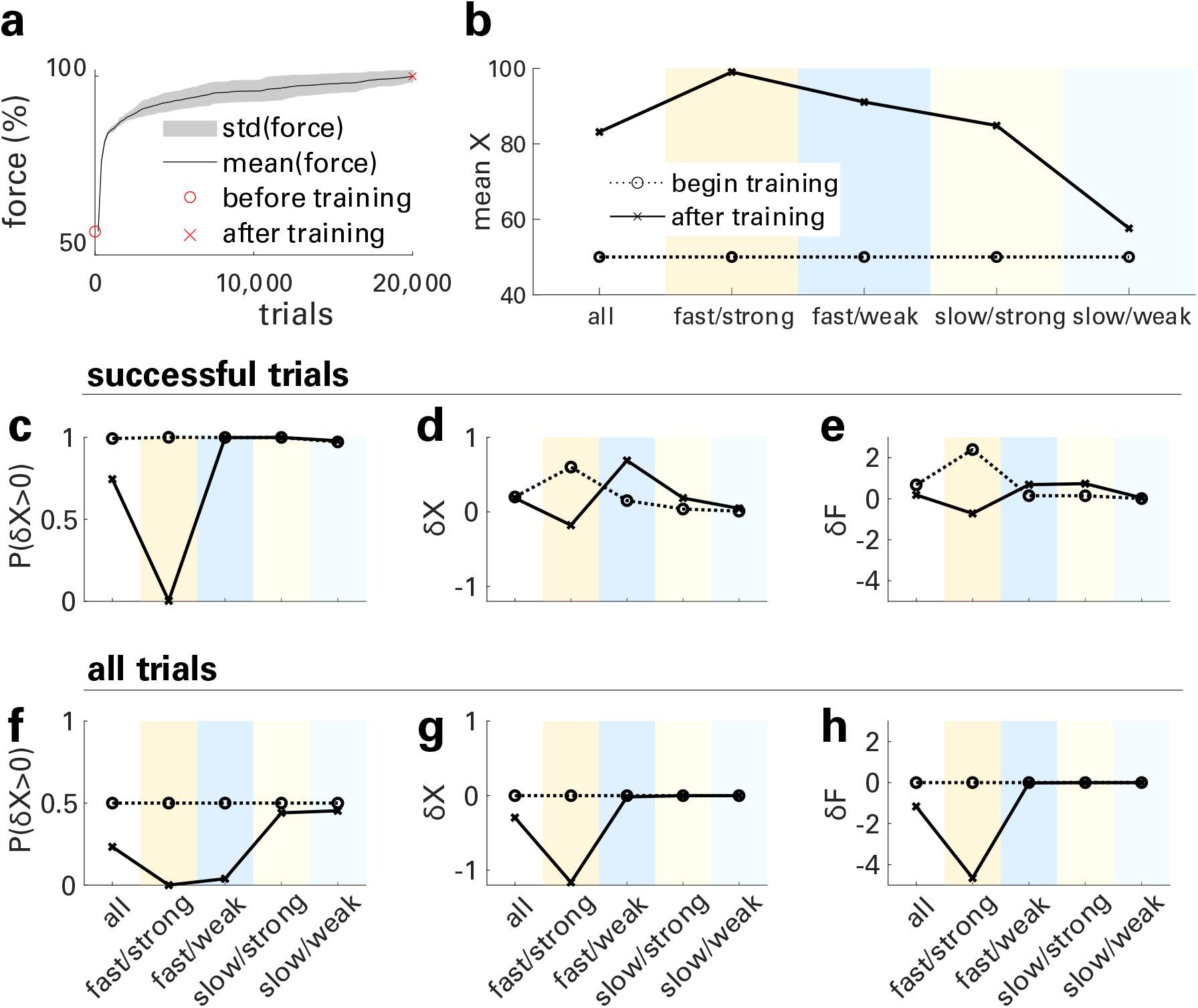
Network effect statistics. **a**, Total force generation (as percentage of maximum) over 20,000 training trials. **b-h**, Neuronal population measures for trials at the beginning of training (dotted lines) and after 20,000 trials (solid lines), calculated by a Monte-Carlo simulation. **b**, Mean activation levels for each population. **c**, Probability that each population’s mean change in activation was positive on a successful trial (i.e., a trial that increased the total force). **d**, Mean change in each population’s activation level for successful trials. This is effectively the rate of optimization. **e**, Each population’s mean contribution to the force change on a successful trial. This indicates which population the model is currently optimizing. **f**, Probability that a population’s mean change in activation was positive on any given trial. **g**, Mean change in a population’s activation level for all trials. **h**, A population’s mean contribution to the force change on all trials.

## Fast but weak neurons optimize before slow but strong neurons

Because the training signal is total force, the network preferentially optimizes neurons by their force change, not their activation change. Late in learning, fast/weak and slow/strong neurons are similar in mean force change per successful trial (Fig. 8e, after training), but fast/weak neurons increase activity more on successful trials, thus optimizing more quickly (Fig. 8d, after training). As these neurons saturate, they also block optimization of the slow/strong neurons, and thus they limit total force recovery.

With this understanding, we can now return to the questions asked at the beginning of this section:

### Why did activity lateralize in the undamaged brain?

The network favors optimization of high-variability/strong-connectivity (fast/strong) neurons, which are located mainly in primary motor cortex (M1) contralateral to the finger movement.

### Why did activity become bilateral after the simulated stroke?

After injury deprived the network of fast/strong neurons, it preferentially optimized fast/weak neurons, which are located mainly in M1 ipsilateral to the finger movement (Fig. 5B).

### Why did targeted plasticity improve force recovery and re-lateralize hemispheric activity?

Standard training blocked optimization of slow neurons. By focusing on secondary motor areas (where slow neurons are mainly located), TNP training overcame the blocking and enabled optimization of the slow neurons, which are stronger contralaterally.

## Discussion

After a stroke, electroencephalography (EEG) and functional magnetic resonance imaging (fMRI) studies reveal substantial reorganization of movement-related cortical activity (Cramer and Crafton, 2006; Wolpaw and Carp, 2006; Yozbatiran and Cramer, 2006; Calautti *et al.*, 2007, 2010; Wu *et al.*, 2010). For example, unilateral movements that are contralateral to the affected hemisphere can elicit bilateral activity (Cramer *et al.*, 1997; Fu *et al.*, 2006; Wu *et al.*, 2010; Rossiter *et al.*, 2014). This loss of hemispheric laterality correlates with decreased motor function. It may reflect a suboptimal compensatory strategy that ultimately limits motor recovery (Cramer and Crafton, 2006; Calautti *et al.*, 2010). The present study uses a computational model to explain these shifts in laterality as arising from the motor system’s stochastic search for neurons that can help drive motorneuronal pools following injury. The model further demonstrates the ability of a new therapeutic strategy, targeted neuroplasticity (TNP), to modify abnormal movement-related cortical activation and to thereby improve motor recovery (Levin *et al.*, 2008).

In traditional rehabilitation, patients simply practice the skills that have been impaired by stroke (e.g., locomotion, reach and grasp, speech). In the model, practicing finger extension causes the neural search process to settle on a suboptimal pattern of activation.

In contrast, the TNP protocol simulated here uses operant conditioning to modify the task-related activation of specific subpopulations of corticospinal (CS) neurons and to thereby enhance the functional recovery of the entire population. In both animals and humans, TNP protocols can target beneficial plasticity to a specific CNS site (e.g., a spinal reflex pathway or a cortical area) by operantly conditioning EMG or EEG features that reflect activity in that site (Chen *et al.*, 2006; Sitaram *et al.*, 2016). This plasticity improves function and enables wider plasticity that further improves function (Thompson *et al.*, 2013; McFarland *et al.*, 2015; Pichiorri *et al.*, 2015). For example, a TNP protocol that operantly conditions a spinal reflex pathway can improve walking in rats (Chen *et al.*, 2006) or people with incomplete spinal cord injuries (Thompson *et al.*, 2013). And a TNP protocol that operantly conditions specific EEG features can enhance functional recovery after stroke (Buch et al., 2008; Norman et al., 2018; Pichiorri et al., 2015; Ramos-Murguialday et al., 2013).

To date, only a handful of studies have attempted to model the mechanisms underlying sensorimotor rehabilitation (Reinkensmeyer *et al.*, 2016; Sedda *et al.*, 2018); none has modeled a TNP protocol. In this paper, we do this by building on an approach that employed a simplified corticospinal (CS) neural network with inherent stochastic noise to simulate finger movement recovery after stroke (Reinkensmeyer *et al.*, 2012). The model used a biologically plausible reinforcement learning (operant conditioning) algorithm to optimize CS activation patterns over repeated motor practice. This network reproduced major clinically observed features of motor recovery after stroke, including exponential recovery and latent residual capacity. A subsequent paper extended this model to simulate multiple limbs and explore the effects of strength and coordination training after neurological injury (Norman, Lobo-Prat, & Reinkensmeyer, 2017). Here, we extend the model using biologically plausible neuronal population parameters and employ it to predict the results of combining a standard rehabilitation protocol with a TNP protocol.

The computational model presented here sheds light on the mechanisms behind the reorganization of neural activity after stroke. It supports the conclusion that, after a stroke damages cortex, traditional rehabilitation methods fail to optimize cortical reorganization; this limits motor recovery. Furthermore, it supports the hypothesis that appropriate TNP interventions can improve cortical reorganization and enhance motor recovery. We found that Scenario 2, which simulated traditional rehabilitation, left the network with residual (i.e., unaccessed) capacity for motor recovery. Scenario 3, which incorporated TNP training, was able to access much of this residual capacity.

While mounting evidence suggests TNP interventions affect cortical reorganization, it is not clear how to maximize their beneficial effects on functional recovery (Cramer *et al.*, 2011). Our reductionist model provides important, and surprising, insight. It shows that targeting neurons that are not easily accessed by regular motor practice is particularly beneficial. With conventional training, these neurons are blocked from full optimization. TNP circumvents this blocking and thereby increases total force. Furthermore, the model results indicate the optimal dosage and target of TNP training. The following sections compare the model predictions regarding network reorganization with the results of imaging studies, summarize TNP principles derived from the model, and consider the study’s limitations and the most promising and important directions for further research.

### Cortical organization before and after cortical injury

Extensive EEG and fMRI data indicate that activation of contralateral primary motor areas normally precedes and accompanies motor actions (Kim *et al.*, 1993; Pfurtscheller and Lopes da Silva, 1999; Grefkes and Fink, 2011). In accord with these experimental data, our model, applied prior to stroke, optimized mainly the high-variability, high-connectivity (i.e., fast/strong) neurons concentrated in contralateral primary motor cortex. Thus, learning a right-hand task optimized activation of neurons in the primary motor cortex of the left hemisphere.

After the simulated stroke destroyed primary motor cortical neurons, the model exhibited a profound but suboptimal reorganization of network recruitment. This phenomenon is prominent in clinical data (Zemke *et al.*, 2003; Yozbatiran and Cramer, 2006; Grefkes and Fink, 2011). Injuries to sensorimotor regions often result in extension of motor representation into perilesional regions (Weiller *et al.*, 1993; Nudo *et al.*, 1996; Muellbacher *et al.*, 2002; Cramer and Crafton, 2006). Another effect is increased activation in the uninjured hemisphere (Chollet *et al.*, 1991; Murase *et al.*, 2004); this is especially prominent for unilateral movements of the affected limb (Cramer *et al.*, 1997; Wu *et al.*, 2010; Grefkes and Fink, 2011). In accord with these clinical data, after the stroke, the model produced diffuse perilesional activation in the lesioned hemisphere and a prominent increase in activation of the uninjured hemisphere.

The model provides insight into the possible mechanisms of this suboptimal reorganization. For a network that uses stochastic search to optimize neuronal activation, the trial-to-trial variability of each neuron, and, secondarily, its connectivity to the motoneuronal pool, determines the rate at which the network is able to recruit that neuron (i.e., to increase its contribution to the total finger extension force). Once the most quickly optimized neurons (e.g., those in contralateral primary motor areas) are saturated, they can only remain saturated or incrementally decrease their activity as they experience trial-to-trial variability. After stroke, these incremental decreases in their activity block optimization of other neurons. While the shift toward bilateral activation does increase force, it is suboptimal because it blocks recruitment of high-connectivity (but more slowly recruited) neurons in other areas that could better enhance total force. This leaves a substantial residual (i.e., unused) capacity for further recovery of force.

TNP training partially re-lateralized hemispheric activity and improved force recovery. Such re-lateralized activity has also been found to accompany better functional outcome in the stroke recovery literature (Dong *et al.*, 2006). The model provides insight into how TNP can improve motor recovery. As discussed above, neurons in secondary motor areas with the potential to contribute to force production may be blocked from optimization by neurons in primary motor cortical areas with higher trial-to-trial variability (i.e., neurons that are rapidly recruited). A targeted intervention can remove this blocking effect by increasing or reducing the roles of specific cortical areas in determining the teaching signal (i.e., in determining whether a trial is successful and thus updates the activation levels of all the CS neurons). The TNP trials of the present study removed the block by ignoring the high-connectivity/high variability neurons concentrated in primary motor areas and deriving the teaching signal solely from the low-variability neurons concentrated in secondary motor areas.

In summary, the model replicated the patterns of network organization that are found in people before and after a unilateral cortical injury (e.g., a stroke). Furthermore, it identified a mechanism that can account for both the normal (pre-stroke) and abnormal (post-stroke) patterns: the dynamics of a stochastic search. With a stochastic search, the final topography of activation reflects the topographies of neuronal variability and connectivity across the two hemispheres. Stroke changes those topographies, and thereby changes the results of the stochastic search. TNP training can further modify these results. Used appropriately, it can thereby improve functional recovery.

### Computational Principles of Targeted Neuroplasticity

These results provide a rationale for using TNP training to enhance neuronal recruitment after injury and thereby improve recovery of motor function. The first principle is that interspersing TNP trials that enable recruitment of under-used neuronal populations with standard trials can access residual force capacity inaccessible to standard training alone. This principle is consistent with animal and human evidence that appropriately-designed TNP therapy can induce widespread adaptive plasticity leading to network reorganization and enhanced motor function (Buch et al., 2008; Chen et al., 2006, 2014; Cramer et al., 2011; Christopher deCharms, 2008; Norman et al., 2018; Pichiorri et al., 2015; Ramos-Murguialday et al., 2013; Thompson et al., 2013; Thompson & Wolpaw, 2015; Wolpaw 2018). In these TNP studies, real-time neuroimaging (i.e., EEG or fMRI) or measurement of a key physiological parameter (e.g., an H-reflex) provides the teaching signal. Stated in behavioral terminology, this teaching signal operantly conditions the person to modify key aspects of CNS activity (e.g., the activation levels of neurons in a specific cortical area).

A second principle based on the model is that, as TNP dose (i.e., TNP trials as % of all trials) increases, overall motor recovery increases up to a maximum and then declines. For the model presented here, maximum recovery occurs with a dose of ~10%; this value is not affected by the number of neurons in the model, injury size, or neuronal parameter distributions. This optimum exists because standard trials alone leave a large latent residual capacity; and TNP trials alone leave the majority of neurons untrained. A proper balance between them is essential. This suggests that studies such as (Buch *et al.*, 2008), which provided only TNP trials, might achieve still better results by interspersing TNP trials with standard, non-targeted trials.

### Model limitations and future directions

Our model greatly oversimplifies cortical control of movement and the potential consequences of cortical stroke. It reduces the many pathways that interconnect the cortex and other brain areas with the spinal cord and the motoneurons to a group of weighted connection. It also ignores the numerous and poorly understood effects of a stroke (e.g., the impact of impairment of the somatosensory input that guides and maintains motor performance). Furthermore, human CS neurons reside in multiple cortical areas (i.e., primary motor, supplementary motor, premotor, somatosensory, cingulate, and parietal). The model simplifies this complex reality into high and low-variability neurons that represent primary and secondary motor areas, respectively. Nevertheless, the model’s results are consistent with clinical data. It displays, and helps to explain, phenomena that underlie both normal motor learning and rehabilitation after stroke (Reinkensmeyer *et al.*, 2016).

At the same time, the model’s value might increase with additional complexity. For example, a particularly valuable addition could be capacity for changes in neuronal connectivities to the motoneuron pools; in addition to training neural activation patterns, the network would also train the strengths of individual neuronal connections to the motoneurons. This addition would be particularly relevant given the ongoing discussion within the motor learning literature concerning the multiple learning mechanisms that drive short-term adaptation and long-term learning (Wolpaw & O’Keefe, 1984; Thompson et al., 2009; Zhou, Tien, Ravikumar, & Chase, 2019).

Although this model applies well to corticospinal control of a muscle, it might not apply as well to other rehabilitation problems. To be useful for a given problem, a model must simulate the motor behavior to be restored and the CNS mechanisms that underlie the behavior and its impairment by injury or disease. A variety of existing or conceivable TNP protocols target plasticity related to specific temporal, kinematic, physiological, or anatomical components of a variety of important sensorimotor behaviors. For example, a TNP protocol can target plasticity: in individual joint control during complex arm movements (e.g., (Klein *et al.*, 2012)); in responses to perturbations during movement (e.g., (Krebs, Hogan, Aisen, & Volpe, 1998)); in physiological measures of activity in key neuronal pathways (e.g., (Thompson *et al.*, 2013)); or by using precisely paired stimuli to change a key CNS site (e.g., (Bunday *et al.*, 2018)). As CNS imaging and stimulation technologies continue to improve, the variety and precision of TNP protocols should increase. Hopefully, parallel advances in computational modeling will enable models to facilitate and enhance these advances.

Finally, clinical TNP protocols might incorporate methods suggested by modeling without needing additional technology (e.g., to record and analyze EEG or EMG signals). For example, the present study shows that incorporating trials that remove the influence of fast/strong CS neurons can access recovery capacity that is inaccessible to standard training alone. As with standard training in our model, traditional rehabilitation for finger extension is likely to improve the performance of stronger, but not weaker, motor units. Stronger motor units are easily fatigued, while weaker units are not (Hudspeth *et al.*, 2013). Thus, a protocol that required prolonged maintenance of position against a steadystate torque could eliminate the contributions of stronger motor units and focus training on the weaker units; it might thereby access otherwise inaccessible capacity for force recovery.

In summary, computational models can accelerate the creation and development of novel neurorehabilitation therapies. In contrast to clinical studies, which are generally demanding and time-consuming for both patients and investigators, modeling allows rapid assessment of many different designs and parameter selections. Properly applied, modeling could guide selection of the most promising protocols for actual clinical study and could thereby enable efficient and effective realization of new therapies.

## Acknowledgments

We thank Dr. Peter Brunner for valuable comments on an earlier version of this manuscript.

The National Center for Adaptive Neurotechnologies (NCAN) is a Biomedical Technology Resource Center (BTRC) of the National Institute of Biomedical Imaging and Bioengineering (NIBIB) of the National Institutes of Health (NIH).

## Data availability

URL will be made available at time of publication

## Supplemental Materials

### Selection of neuronal parameter distributions

In preliminary tests, we sampled model parameters (i.e., synaptic connectivity, neuron variability, initial activation patterns (i.e., *X_0_*)) from uniform, normal, or lognormal distributions of varying magnitudes. The three distributions yielded comparable results. We elected to use lognormal distributions because they best reflect physiological and anatomical data. Functional and structural parameters in the brain, including synaptic connectivity and firing rates, are strongly skewed with a heavy tail; they are closely approximated by lognormal distributions (Buzsáki and Mizuseki, 2014).

### Characterizing the learning curve with residual capacity

From a practical clinical perspective, the key property of the model’s results is the residual capacity for force production that remains after training. That is, all three scenarios produced learning curves of force production that did not reach their full potential, even over 20,000 trials, but rather appeared to approach a suboptimal asymptote (Fig. 3). This result is consistent with clinical evidence of actual force recovery after injury (Barreca *et al.*, 2003; Page *et al.*, 2004; Ada *et al.*, 2006; French *et al.*, 2007; Stinear *et al.*, 2007; Kwakkel *et al.*, 2008). In fact, the force profile does continue to increase (similar to a power curve (Newell and Rosenbloom, 1981)), but at an increasingly slow rate. A power curve never saturates. We simulated 5.5 million trials; the network recovers ever-smaller amounts of force at higher amounts of practice. Given infinite time, it approaches an asymptote due to the nonlinear maximum firing rate function *g*, described in Eq. 1.

Thus, the learning curves described in Fig. 3 do not obey a power curve at large time scales. On the other hand, a single exponential function cannot accurately describe both: 1) the initially fast learning of the network; and 2) its latent residual capacity at even moderately large numbers of trials. Force production over time appears to be the sum of fast and slow exponential curves. This double-exponential learning curve can be defined:

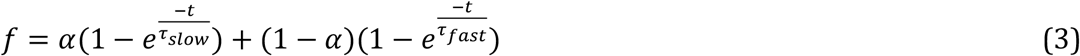

where τ represents the time constant of each curve, *α* is a proportionality constant, and the compound curve sums to 1 as *t* → ∞. This equation, a mixed-exponential learning curve first shown in human motor learning (Newell and Rosenbloom, 1981), adequately describes learning in the model presented here and in (Reinkensmeyer *et al.*, 2012); it includes the compound curvature and the latent residual capacity.

